# dCas9-metabolic enzyme fusions modulate global and locus-specific gene expression

**DOI:** 10.64898/2026.05.10.724078

**Authors:** Kellen V. Biesbrock, Spencer A. Haws, Harshini Cormaty, Rupa Sridharan, John M. Denu

**Author notes:** **Correspondence** John M. Denu, PhD, Professor of Biomolecular Chemistry, University of Wisconsin-Madison, 330 N Orchard St., Madison, WI 53715. These authors contributed equally (Kellen V. Biesbrock, Spencer A. Haws).

## Abstract

Central metabolites function as essential co-substrates for chromatin-modifying enzymes, directly linking cellular metabolism to chromatin regulation. Accordingly, whole-cell fluctuations in co-substrate availabilities have been shown to promote diverse phenotypes through chromatin-dependent mechanisms. There is emerging evidence that metabolic enzymes producing co-substrates for chromatin modifying enzymes can exist in the nucleus, suggesting that nucleus-specific metabolite availability regulates chromatin state. Here, we developed CRISPRm (CRISPR metabolite) to assess how nucleus-specific metabolic perturbations influence chromatin function. Five dCas9-metabolic enzyme fusions (*i.e*., dCas9-ACSS2, -NMNAT1, -MAT2A, -GDH, and -AHCY) were used to modulate nuclear levels of essential co-substrates involved in histone (de)acetylation and (de)methylation reactions. Transient expression of all dCas9 fusions in HEK293T cells induced distinct global changes in gene expression patterns, with dCas9-ACSS2 (acetyl-CoA producing) and NMNAT1 (NAD^+^ producing) eliciting large opposing changes in gene expression, suggesting transcriptional responses to nuclear acetyl-CoA and NAD^+^ production may be directly facilitated by acetylation or deacetylation reactions, respectively. Targeting dCas9-ACSS2 and -NMNAT1 to promoters of select candidate genes revealed enhanced transcriptional modulation. dCas9-ACSS2 upregulated, and dCas9-NMNAT1 downregulated genes showed basal enrichment of H3K9ac, H3K18ac, H3K27ac, H3K4me3, and p300, suggesting these genomic loci reside within epigenetic environments susceptible to fluctuations in acetyl-CoA and NAD^+^ availability. Of significant genes altered, dCas9-MAT2A (SAM producing) increased expression of 72% whereas dCAS9-GDH (alpha-ketoglutarate producing) decreased expression of 79%. Surprisingly, dCAS9-AHCY (SAH hydrolysis) led to down-regulation of shared genes up-regulated by dCas9-MAT2A. The observations amongst the methylation-specific enzymes revealed unexpected and unique gene-regulatory sensitivities to SAM, SAH and alpha-ketoglutarate. Together, these results demonstrate the utility of CRISPRm in studying nuclear metabolic regulation of transcription and provide strong evidence that perturbations in nuclear co-substrates do not lead to a large mass- action changes in chromatin acetylation/methylation but rather to modulation of select chromatin-modifying enzymes with targeted transcription responses.

**Highlights:**

- CRISPRm is a novel, modular dCas9-effector platform that enables interrogation of the metabolism-epigenome axis
- dCas9-ACSS2, -NMNAT1, -MAT2A, -AHCY, and -GDH induce distinct transcriptional programs.
- Targeting CRISPRm to promoters enhances transcriptional responses.
- dCas9-ACSS2 and -NMNAT1 sensitive gene promoters exhibit unique enrichment of chromatin features, including H3K9ac, H3K18ac, H3K27ac, H3K4me3 and p300.

## Introduction

Metabolism and epigenetics are fundamentally linked. The histone-modifying enzymes that create and maintain epigenetic signatures require central metabolites as co-substrates to perform their catalytic activities. Acyl-CoAs are used by histone acetyltransferases, nicotinamide adenine dinucleotide (NAD^+^) by sirtuin deacetylases, *S*-adenosyl methionine (SAM) by histone methyltransferases, and alpha-ketoglutarate (αKG) by JmjC domain-containing demethylases. Therefore, varying intracellular levels of these key metabolites can directly impact chromatin states. For example, modulation of intracellular metabolite pools, including SAM (1–6), NAD^+^ (7), αKG (8, 9), and acyl-CoAs (10–13), has been shown to alter the corresponding histone modifications. However, these studies rely on whole-cell metabolic perturbations, making it difficult to attribute changes in chromatin state to direct effects of nuclear metabolite availability.

In recent years, growing attention has been paid to the compartmentalization of metabolism in the nucleus (14, 15). Distinct nuclear pools of acyl-CoA metabolites, including propionyl-CoA enrichment, have been described (15). Historically, the nucleus and cytosol were assumed to be a single, continuous metabolic compartment based on nuclear pore complex size cut-offs. However, nuclear and cytosolic metabolite composition can diverge under certain conditions. One mechanism of this divergence is the translocation of metabolic enzymes to the nucleus. ATP citrate lyase (ACLY), acyl-CoA synthetase short-chain family member 2 (ACSS2), and pyruvate dehydrogenase (PDH) complex have been shown to translocate to the nucleus and increase histone acetylation in response to DNA damage and metabolic stress, respectively (13, 16, 17). Disruption of NAD^+^-producing enzymes, nicotinamide phosphoribosyltransferase (NAMPT) and nicotinamide mononucleotide adenylyltransferase1 (NMNAT1), decreased intracellular NAD^+^ synthesis and led to decreases in histone and non-histone protein acetylation (18, 19). In addition to its canonical role in mitochondrial glutamine-based anaplerosis, glutamate dehydrogenase (GDH) was reported to produce α-KG in the nucleus and stimulates Tet3 DNA demethylation in neurons (20). Nuclear localization of adenosylhomocysteinase (AHCY) during early mammalian development coincides with reduced methylation and permissive chromatin signatures at the TSS of proliferation-associated genes (21). Importantly, much of this regulation is directed at specific gene loci via interactions with transcription factors that target specific gene sets. ACSS2 nuclear translocation promotes macroautophagy via interacting with TFEB and enables the generation of Acetyl-CoA locally at promoter regions of target lysosomal and autophagosomal genes for H3 acetylation (16). Similarly, AHCY forms a complex with BMAL and is required for rhythmic regulation of H3K4me3 at circadian-controlled genes and methionine adenosyltransferase II (MAT2A) forms a complex with MAFK and represses MARE (22, 23). Despite the recognition that distinct nuclear pools exist, the ability to specifically produce nuclear metabolites, and thus test functional consequences of nuclear metabolite production, remains limited. Moreover, given the importance of targeted metabolite production for fundamental cellular processes, a tool capable of targeting metabolic enzymes to specific genomic loci would prove useful in studying the metabolism-- epigenome axis. This motivated the development of a tool that enables nuclear compartment perturbation, with the option to target specific genomic loci of interest.

The dCas9 platform has proven useful for studying transcriptional and epigenetic regulation. Various effector domains have been fused with dCas9 to activate or repress transcription, such as the transactivator domain of the Herpes Simplex viral protein 16 (VP16) and KRAB domain of Kox1 (24–26). Many studies have validated the ability to modulate site-specific chromatin signatures and regulate the expression of genes by fusing dCas9 to different histone-modifying enzymes. For example, dCas9 fused to the catalytic core of p300 increased transcription and enriched H3K27ac at target genes when targeted to promoters and enhancers (27). A reasonable extension of this platform would suggest that any enzyme whose catalytic activity or effector recruitment has been shown to have consequential effects on chromatin and gene expression could be utilized to investigate the role of global vs. targeted delivery of essential co-substrates for chromatin modification.

Here, we developed and investigated five dCas9-metabolic enzyme fusion constructs, referred to as CRISPRm, utilizing metabolic enzymes known to localize to the nucleus and to influence histone modification. To target both major histone modification pathways of acetylation and methylation, we selected ACSS2 and NMNAT1 as modulators of nuclear acetylation, and MAT2A, AHCY, and GDH as modulators of nuclear methylation. We hypothesized that dCas9-metabolic enzyme fusions would modulate distinct transcriptional programs defined by sensitivity to nuclear metabolic enzyme activity and enable identification of the chromatin features underlying those differential responses. The CRISPRm system enabled the systematic evaluation of nuclear metabolic enzymes, revealing enzyme-specific transcriptional programs that may arise from metabolite-dependent chromatin modification, protein-protein interactions, or other mechanisms. Additionally, the system facilitated the identification of local chromatin features associated with CRISPRm-sensitive genes. Furthermore, the capacity to direct fusion constructs to specific genomic loci enabled precise examination of the transcriptional effects of localized metabolic enzymes at gene promoters. Overall, CRISPRm provides a platform for probing the diverse mechanisms by which nuclear metabolic enzymes influence gene expression.

## Results

### CRISPRm is an effective tool for manipulating cellular metabolism

We engineered each metabolic enzyme (ACSS2, NMNAT1, MAT2A, AHCY, and GDH) fusion construct from an HA-dCas9-mCherry backbone, which was generated by adding an SV40-driven mCherry reporter, an N-terminal 2X HA tag, and removing the C-terminal KRAB domain from dCas9-KRAB (28). To confirm expression of dCas9-fusion constructs, HEK293T lysates were analyzed by Western blot using an anti-HA antibody at 48 hours post-transfection (Fig. 1A). All five dCas9-fusion constructs produced bands at the expected molecular weight, consistent with full-length translation. Next, HEK293T cells underwent flow cytometry for mCherry fluorescence and DAPI exclusion to further evaluate the relative transfection efficiency and cell viability, respectively, 48 hours after transfection of each construct, (Fig. 1B). All dCas9 constructs exhibited transfection efficiency comparable to WT-dCas9, except for dCas9-ACSS2, which was approximately 10% higher, at ∼70%, confirming similar rates of plasmid uptake and viability across conditions. Whole-cell metabolite extraction and LC-MS metabolite quantification were performed 48 hours post-transfection to assess the ability of the dCas9-metabolic fusion proteins to modulate the levels of target metabolic substrates or products (Fig. 1C-H). dCas9-ACSS2, -MAT2A, and -AHCY induced whole-cell changes in Acyl-CoAs, SAM, and SAH, respectively. dCas9-ACSS2 increased both acetyl-CoA and propionyl-CoA relative to WT-dCas9 (Fig. 1 C,D). dCas9-MAT2A increased SAM abundance and dCas9-AHCY decreased SAH abundance relative to WT-dCas9 (Fig. 1F, G), while dCas9-NMNAT1 and -GDH expression showed no significant changes in whole-cell levels of NAD^+^ or αKG, respectively. The absence of detectable whole-cell changes in NAD^+^ and αKG was not unexpected and does not preclude hypothesized nuclear perturbation, given that whole-cell metabolomics does not reliably reflect nuclear metabolite concentrations (15). Moreover, steady-state metabolomics cannot measure shifts in flux of highly interconnected metabolites, such as αKG. Together, these results indicate that CRISPRm constructs are expressed in the nucleus and lead to significant alterations in the steady-state levels of the corresponding target metabolites for dCas9-ACSS2, -MAT2A and -AHCY.

**Figure 1:**
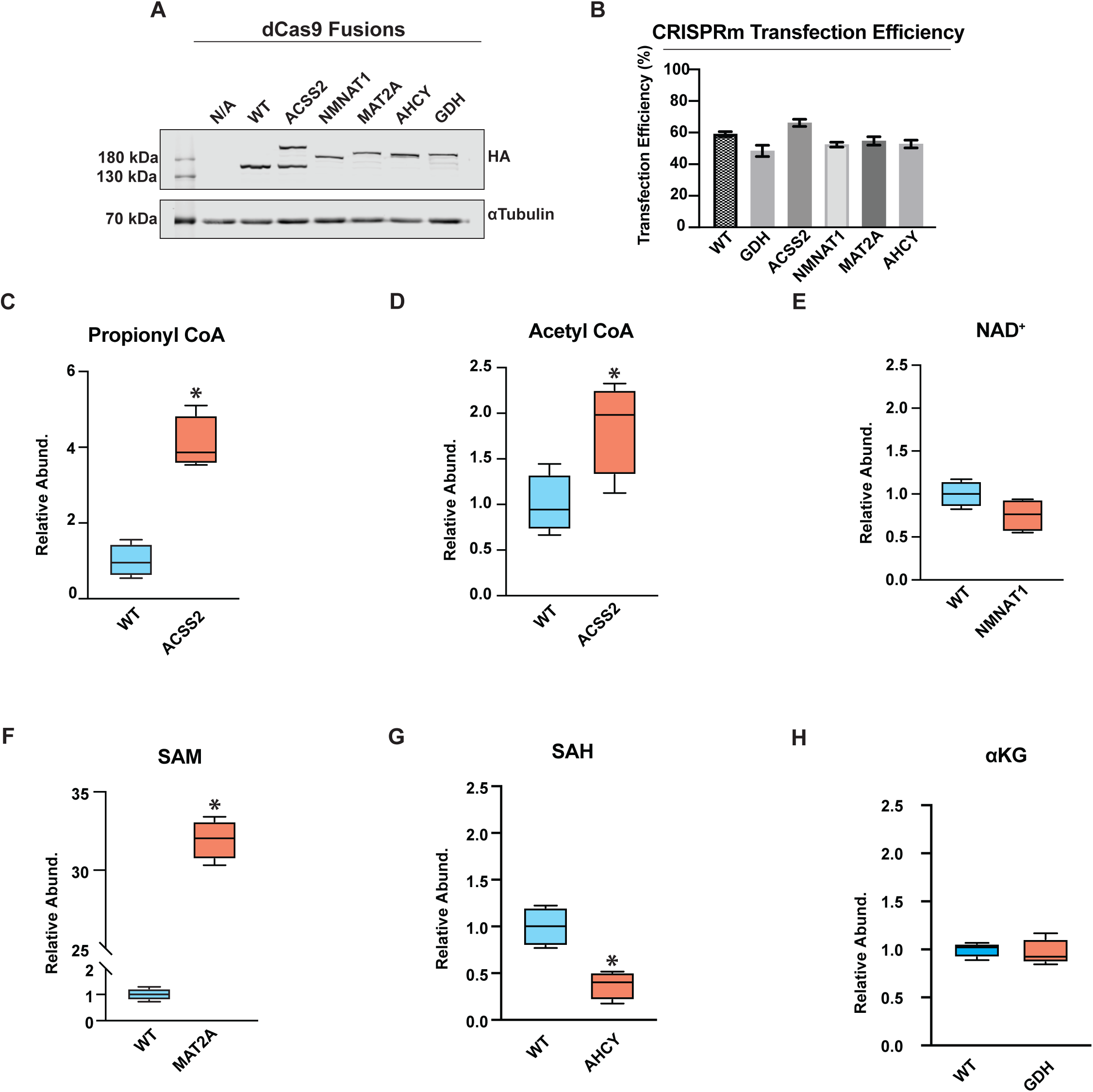
CRISPRm validation experiments. (*A*) Anti-HA Western blot of whole-cell lysates from HEK293T cells transfected with WT-dCas9, dCas9-ACSS2, -NMNAT1, -MAT2A, -AHCY, and -GDH. Alpha-tubulin is shown as a loading control. (B) Transfection efficiency in HEK293T cells, calculated as the percentage of DAPI-negative (live) cells that are mCherry-positive. Bars show mean ± SD; *n* = 6 biological replicates per construct (dCas9-GDH: n = 5). Quantification performed by flow cytometry. (C-H) Targeted LC-MS quantification of CRISPRm-relevant metabolites in HEK293T cells expressing the indicated constructs: (C) propionyl-CoA, (D) acetyl-CoA, (E) NAD⁺, (F) SAM, (G) SAH, and (H) αKG. Metabolite levels are normalized to total protein (BCA) and shown relative to WT-dCas9. Boxplots shown; *n* = 4 biological replicates. Statistical comparison vs. WT-dCas9 by Student’s two-tailed *t*-test; * p < 0.05.

### CRISPRm fusion constructs elicit global changes to transcriptional programs

To assess the transcriptional response to nuclear-localized metabolic enzymes, we performed bulk RNA-sequencing 48 hours post-transfection for all five CRISPRm constructs. Principal Component Analysis (PCA) of log-transformed gene counts per million values (Fig. 2A) revealed distinct groupings among all constructs compared with WT-dCas9, supporting that these fusion constructs effectively modulated global gene expression relative to dCas9-WT controls. Moreover, each grouping was distinct among the five dCas9-metabolic enzyme fusion constructs, demonstrating that the transcriptional responses were specific to the identity of the nuclear enzyme rather than to common secondary signaling or compensatory mechanisms. In support of transcriptional regulation in response to enzyme identity, 68.9% (n = 1809), 45.3% (n = 1065), 52.5% (711), 23.1% (n = 540), and 24.7% (n = 648) of differentially expressed (DE) genes (|log2FC| > 0.25 & padj < 0.05) were unique to dCas9-ACSS2, -NMNAT1, -MAT2A, -AHCY, and -GDH, respectively (Fig. 2B). Moreover, only 0.2% (n = 3500) of differentially expressed (DE) genes were common to all 5 groups. Globally, dCas9-ACSS2 and -MAT2A predominantly upregulated transcripts while dCas9-NMNAT1 and -GDH predominantly downregulated transcripts (Fig. 2D, G, E, I). Globally, dCas9-AHCY showed near-equal activation and repression of transcripts (Fig. 2H). Together, these data support that CRISPRm is sufficient to induce enzyme-specific transcriptional responses.

**Figure 2:**
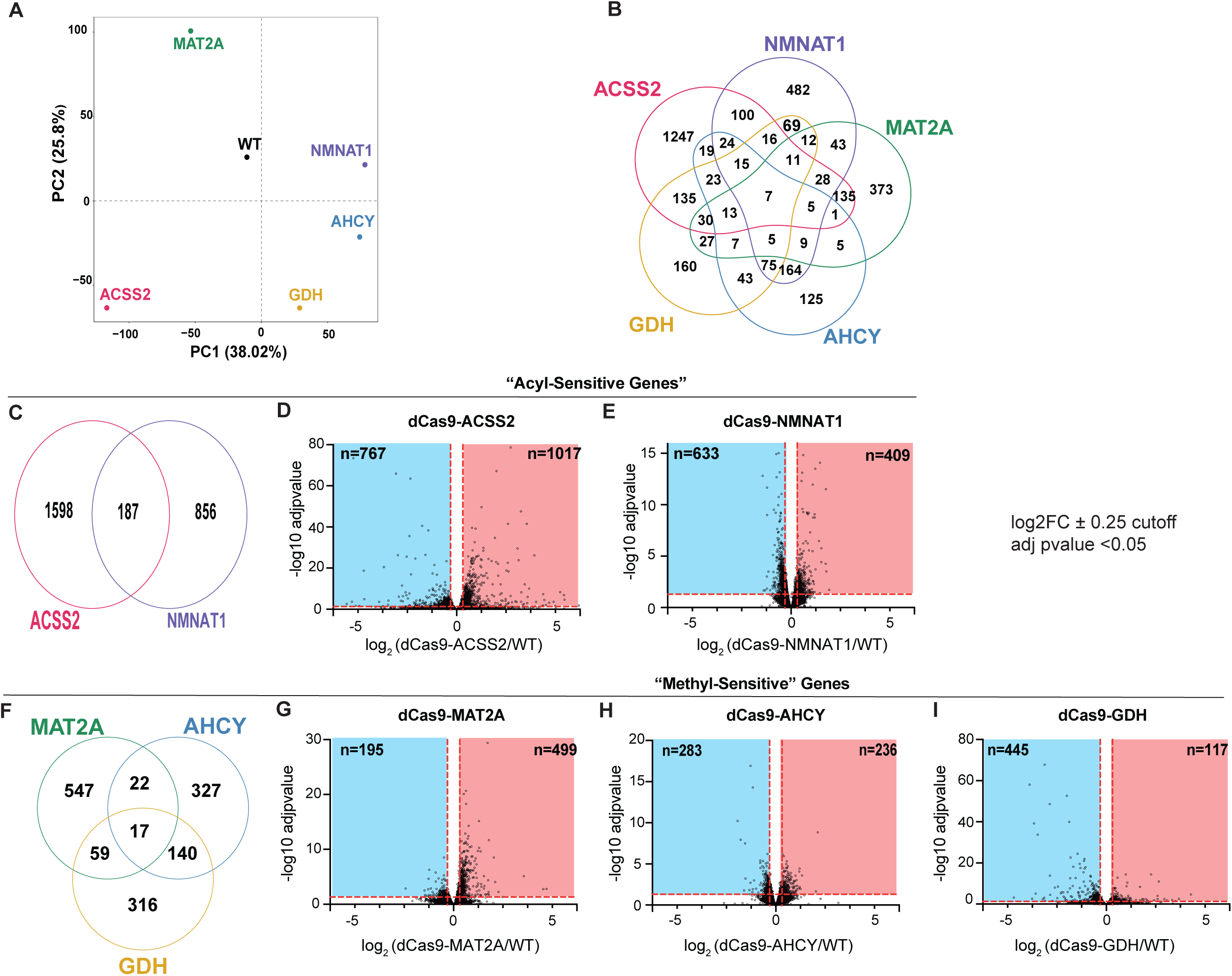
Global transcriptional responses to CRISPRm perturbations. (*A*) PCA analysis of log transformed gene CPM values. (*B*) Venn diagram of differentially expressed genes of all CRISPRm constructs. *(C)* Venn diagram of differentially expressed genes at p-adj.<0.05 and |log2FC| > 0.25 for dCas9-ACSS2 and -NMNAT1 relative to WT-dCas9. (D-E) Volcano plots depicting log2FC and -log10 adj. *p*-values for all identified genes as differentially expressed between WT and dCas9-ACSS2 and -NMNAT1, respectively. Horizontal and vertical markers indicate threshold for statistical significance (p-adj. < 0.05, |log2FC| > 0.25). *(F)* Venn diagram of differentially expressed genes at p-adj.<0.05 and |log2FC| > 0.25 for dCas9-MAT2A, -AHCY, and -GDH relative to WT. (G-I) Volcano plots depicting log2FC and -log10 adj. *p*-values for all identified genes as differentially expressed between WT and dCas9-MAT2A, -AHCY, and -GDH, respectively. Horizontal and vertical markers indicate threshold for statistical significance (p-adj. < 0.05, |log2FC| > 0.25).

To determine the transcriptional programs regulated by the different CRISPRm constructs, gene set enrichment analysis (GSEA) was performed using Gene Ontology Biological Process (GO-BP) functional annotations for the complete ranked gene list from each construct relative to WT-dCas9. In addition, to prevent bias in transcriptional program identification, we computed pairwise semantic similarity of the enriched terms and visualized the top 50 enriched terms, ranked by adjusted p-value, as tree plots to identify biological processes most regulated by each CRISPRm construct (Fig S2A-E). GSEA of dCas9-ACSS2 genes identified 340 significantly (padj. < 0.05) enriched GO-BP gene sets, including positive enrichment of autophagy (*e.g.,* macroautophagy), and negative enrichment of RNA processing (*e.g.,* rRNA processing) (Fig. S3A, B). dCas9-NMNAT1 genes were enriched for 393 GO-BP gene sets, including positive enrichment of nucleotide biosynthesis (*e.g*., nucleotide metabolic process) and electron transport chain (*e.g*., ATP synthesis coupled electron transport), and negative enrichment of chromatin organization (e.g., heterochromatin formation) and RNA catabolism (e.g., RNA destabilization) (Fig. S3A, C). These results highlight the capability of CRISPRm to recapitulate known transcriptional responses of nuclear-localized ACSS2 and NMNAT1, including promoting autophagy and opposing rRNA transcription, respectively (16, 29). dCas9-NMNAT1’s ability to induce transcriptional programs mirroring NAD^+^-sensitive programs suggests nuclear enrichment of NAD^+^ despite constant whole-cell levels. To identify candidate transcriptional programs sensitive to nuclear acetylation levels, significantly enriched gene sets were compared between analyses of dCas9-ACSS2 and -NMNAT1. Of 64 shared GO:BP terms, 61 exhibited reciprocal enrichment, suggesting transcriptional regulation in response to (de)acetylation (Fig. S3A,D). These terms were primarily related to cell cycle regulation *(e.g.,* mitotic cytokinesis and cell cycle phase transition), consistent with the opposing functions of acetylation and deacetylation in cell cycle progression (30–32). Additionally, there were a GO:BP terms related to ribosome assembly and translation (e.g., ribonucleoprotein complex biogenesis), which recapitulate previously described ribosomal RNA and protein gene regulation by acetylation via Ihf1 and H3K9 acetylation (33, 34).

For the methylation-modulating constructs, GSEA identified 563, 342, and 303 significantly enriched GO-BP terms for dCas9-MAT2A, -AHCY, and -GDH, respectively (Fig. S3E-H). dCas9-MAT2A showed positive enrichment for RNA processing pathways (e.g., mRNA splicing via the spliceosome) and negative enrichment for developmental programs (e.g., renal system development) (Fig. S3F). dCas9-AHCY and -GDH both showed positive enrichment of nucleotide biosynthesis (e.g., purine nucleotide biosynthetic process) and oxidative phosphorylation (e.g., aerobic electron transport chain), and negative enrichment of chromatin organization (e.g., chromatin looping) (Fig. S3G, H). Notably, dCas9-AHCY and -GDH show similar GO:BP term enrichment despite being hypothesized to exert opposing effects on nuclear methylation activity. To identify transcriptional programs broadly sensitive to perturbation of nuclear methylation, significantly enriched gene sets were compared across all three constructs, yielding 62 shared GO-BP terms (Fig. S3E). While dCas9-MAT2A and -AHCY are both predicted to promote nuclear methylation through SAM synthesis and SAH hydrolysis, respectively, they exhibited largely opposing effects on shared gene set regulation: dCas9-MAT2A showed predominantly positive enrichment, while dCas9-AHCY showed predominantly negative enrichment (Fig. S3I). dCas9-GDH, which is predicted to promote demethylation through α-KG production, showed exclusively negative enrichment among shared terms with MAT2A (Fig. S3J). Similarly, the shared 62 GO:BP terms showed concordant enrichment between dCas9-GDH and -AHCY (Fig. S3K). Together, the GSEA data confirms the ability of CRISPRm to induce enzyme-specific transcriptional programs and suggests that the mechanisms reflect unique metabolite regulation of chromatin-modifying enzymes.

To identify candidate genes underlying these pathway-level signatures, we next examined the unique core enrichment genes across all GO:BP terms identified by GSEA for dCas9-ACSS2 and -NMNAT1. Notably, 89.9% (n=1431) and 98.5% (n=1302) of unique core enrichment genes are not DE genes defined by a highly rigorous cutoff , indicating that pathway-level signatures are largely driven by small, statistically insignificant shifts across many genes. To identify candidate genetic loci sensitive to acetylation or methylation, we identified DE genes shared by dCas9-ACSS2 and -NMNAT1, or by any 2 of dCas9-MAT2A, -AHCY, or -GDH datasets (Fig. 2C, F). Of the 187 shared acetyl-sensitive genes, 125 genes were upregulated by dCas9-ACSS2 and downregulated by dCas9-NMNAT1, consistent with activation of genes by acetylation and repression by NAD^+^-dependent deacetylation. Of the 98 shared methyl-sensitive genes between dCas9-MAT2A and dCas9-GDH, 87 genes were upregulated by dCas9-MAT2A and downregulated by dCas9-GDH, suggesting activation of genes by methylation and repression by demethylation. Interestingly, dCas9-AHCY and dCas9-GDH showed near-100% concordance in downregulation of shared genes, despite being hypothesized to have opposing effects on methylation (Fig. 2L). Although we predicted overlap of MAT2A (SAM synthesis) and AHCY (SAH clearance) on common methylation-sensitive targets, dCas9-MAT2A (n = 606) and dCas9-AHCY (n = 467) coregulated only 40 genes, indicating that these enzymes elicit largely distinct transcriptional responses rather than regulating a common set of methylation-sensitive loci. Overrepresentation Analysis (ORA) with GO:BP terms using the subset of reciprocally regulated DE genes for dCas9-ACSS2 and -NMNAT1 did not reveal any significantly enriched terms (padj. < 0.05). Similarly, ORA of shared genes among any two methylation-related CRISPRm constructs revealed no significant GO:BP terms enriched among these genes across all comparisons (Table 1). The absence of significant ORA results, despite GSEA enrichments, further supports that transcriptional programs identified by GSEA are driven by small changes in many GO:BP member genes but do not capture the most sensitive loci to nuclear metabolic perturbation.

### Global Histone PTM profiles are largely resistant to CRISPRm perturbation

A central question underlying metabolite-mediated chromatin regulation is whether altered nuclear metabolite levels drive locus-specific or global changes in histone PTM levels. To investigate the potential role of histone modification in CRISPRm-mediated transcriptional regulation, we performed histone PTM profiling of ∼180 proteoforms following transfection of all five CRISPRm constructs (35). Initially, we examined peptides that showed reciprocal changes in the acetylation-related CRISPRm constructs dCas9-ACSS2 and dCas9-NMNAT1. Remarkably, there was minimal consistent change to canonical histone acetylation sites, including H3K18ac, H3K27ac, H4K16ac, and H3K56ac, suggesting that dCas9-ACSS2 and -NMNAT1 do not lead to changes in global acetylation and NAD^+^-dependent deacetylation of histones (Fig. 3B, S3A, C). Although no individual mark reached statistical significance in both directions, H3K18K23pr showed a reciprocal response that was significantly increased by dCas9-ACSS2 (Fig. 3C, S3A, C). Acetylation at either H3K9 or H3K14 was decreased in both the dCas9-ACSS2 and dCas9-NMNAT1 conditions relative to WT control (Fig. 3D-F). Additionally, dCas9-NMNAT1 led to increased tetra-acetylated H4 (H4K5K8K12K16ac) (Fig. G-I). Beyond acylation, dCas9-ACSS2 and -NMNAT1 led to changes at histone methylation sites (Fig. S2B, D). For example, H3K4me3 was significantly increased in the dCas9-ACSS2 condition relative to WT, consistent with the overall increase in up-regulated gene expression. These results suggest that nuclear-localized metabolic enzymes influence chromatin modifications through mechanisms that extend beyond the simple nuclear enrichment of metabolic co-substrates available to chromatin modifying enzymes. Methylation-related CRISPRm constructs exhibited similar resistance to global changes in histone modification. Histone methylations with well-established roles in transcriptional regulation — including those at H3K4, H3K9, H3K27, H3K36, H3K79, and H4K20 — remained statistically unchanged in response to dCas9-MAT2A, -AHCY, and -GDH (Fig. 3B). However, dCas9-GDH produced noteworthy but non-significant trends in PTMs within H3K27–K36, including decreased H3.3K27me3K36me2 and H3K27me3K36me2 with concurrent increases in H3K27K36un, H3K27me1K36me1, H3.3K36me1, and H3K27me1 (Fig. 3J-M, S3D). This pattern of reduced higher-order methylation states with accumulation of lower methylation states is consistent with enhanced α-ketoglutarate-dependent JmjC domain demethylase activity driven by dCas9-GDH. dCas9-MAT2A showed trends of increased H3K4me3, H3K27me3K36me2, and H3.3K27me3K36me2 while levels of H3K27me3 remained unchanged, suggesting dCas9-MAT2A may preferentially promote modifications associated with activation like H3K4me3 and H3K36me2 (Fig. 3S3D). Similarly, dCas9-AHCY exhibited a trend in decreased H3.3K27me1K36me2 but maintained H3K27me3 levels, consistent with increased methyltransferase activity (Fig. 3L, S3D). To assess whether altering metabolites that impact histone methylation state could result in opposing patterns of histone methylation, we examined peptides reciprocally regulated by dCas9-AHCY and dCas9-MAT2A relative to dCas9-GDH. This analysis identified 14 reciprocally regulated peptides, the majority of which mapped to the H3K27–K40. At this location, dCas9-GDH promoted accumulation of K27 and K36 monomethylation states while dCas9-MAT2A and dCas9-AHCY favored higher methylation states such as K27me3K36me2, further implicating these sites as sensitive to perturbations in nuclear metabolites (Fig. 3K, M, S3D). Together, these findings suggest that most CRISPRm-mediated transcriptional changes are not readily explained by global alterations in histone acetylation or methylation and may instead reflect more targeted, locus-specific changes in histone modifications.

**Figure 3:**
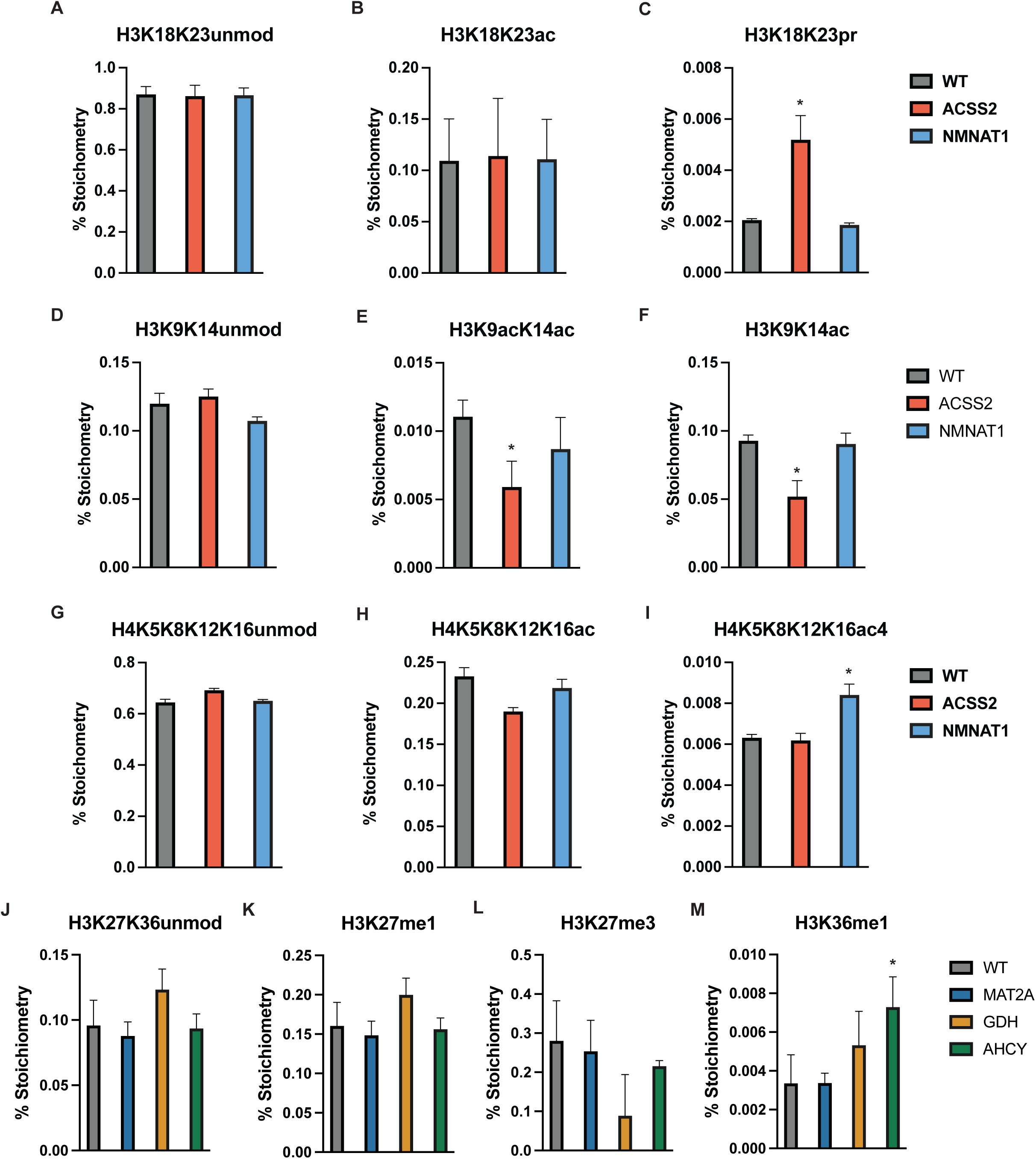
Global histone PTM regulation following CRISPRm expression. (*A-M*) Bar charts depicting deconvoluted stoichiometry percentages of residue-specific histone PTMs. Error bars = SD; N = 4 for WT, N = 4 dCas9-ACSS2, N = 4 dCas9-NMNAT1, N = 4 dCas9-MAT2A, N = 4 dCas9-AHCY, N = 4 dCas9-GDH; Differential modification was assessed using limma with Benjamini–Hochberg correction; * adj. p < 0.05 vs. WT-dCas9.

### CRISPRm acyl- and methyl-sensitive genes exhibit distinct epigenetic modification and modifier enrichment patterns

Given the general lack of global changes observed in histone methylation and acetylation with CRISPRm, yet significant changes in gene expression, we suspected that genes sensitive to CRISPRm were characterized by specific chromatin features. Using publicly available ChIP-seq data, we sought to identify histone acetylations and writers enriched at acetyl-sensitive genes, hypothesizing that specific marks and writers may confer sensitivity to acetyl-metabolite levels via enrichment at these loci. We chose known chromatin marks associated with transcriptional regulation, including H3K9ac, H3K14ac, H3K18ac, H3K27ac, H3K4me1, and H3K4me3. Additionally, we evaluated one histone acetyltransferase, EP300, to gain mechanistic insight into the candidate enzymes responsible for histone acetylation near these loci. We hypothesized that the transcriptional sensitivity of genes to acetylation-related nuclear metabolic enzymes is encoded in the baseline chromatin state of sensitive loci, predicting that acetylation-sensitive genes would be distinguished by enrichment of specific histone modifications rather than a uniform enrichment of all acetylation marks. Consistent with this, acetyl-sensitive genes (upregulated by ACSS2 and downregulated by NMNAT1) showed a relative enrichment in H3K9ac, H3K18ac, H3K27ac, H3K4me3, and EP300 at the promoters of these genes compared to a control group of genes that were sensitive to at least one of the three methyl-metabolite enzymes (Fig. 4). Co-occupancy of EP300 and its associated marks, H3K18ac and H3K27ac, suggests a role in gene regulation of the acetyl-sensitive genes. We identified that 39.5% (n = 1573) of upregulated dCas9-ACSS2 genes and 31.5% (n=1972) downregulated dCas9-NMNAT1 genes were shared, suggesting these 622 genes are sensitive to both acetylation and deacetylation. Moreover, most genes are unique to each construct, suggesting that nuclear-localized dCas9-ACSS2 and dCas9-NMNAT1 regulate gene expression independently of acetylation and NAD+-dependent deacetylation of shared targets. Interestingly, there is comparatively little enrichment of H3K14ac and H3K4me1. KAT7 is required for H3K14ac in human cells (36–38), further suggesting p300’s role in histone acetylation at these sensitive genetic loci. Overall, this data suggests that surrounding histone modifications and chromatin-modifiers at gene loci can confer susceptibility to nuclear metabolite perturbation.

**Figure 4:**
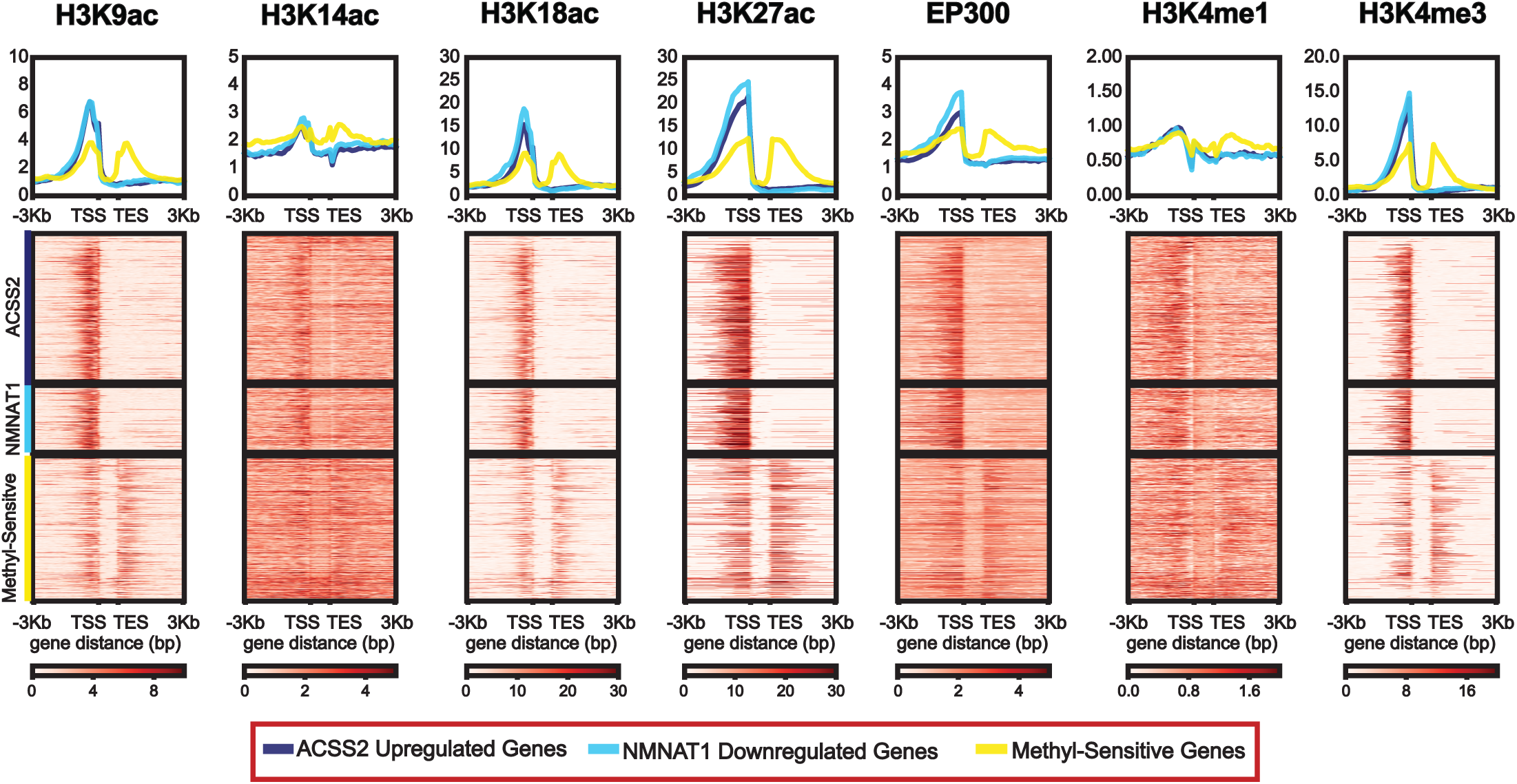
Mapping diverse epigenetic modifications and modifiers to CRISPRm DE genes. Heatmap and composite profiles of H3K9ac, H3K14ac, H3K18ac, H3K27ac, EP300, H3K4me1, and H3K4me3 enrichment around the TSS of ACSS2 upregulated genes, NMNAT1 downregulated genes, and a control set of genes sensitive to any of dCas9-MAT2A, -AHCY, and -GDH termed ‘methyl-sensitive’ genes.

### Locus-specific targeting of NMNAT1 and ACSS2 at CRISPRm acyl-sensitive genes

The CRISPRm metabolic enzymes are required for locus-specific transcriptional regulation and histone modification under different conditions, suggesting loci-specific metabolite production as a mechanism of transcriptional regulation (16, 18, 21, 39–41). To directly test metabolite production, we next sought to study transcriptional regulation induced by CRISPRm targeting specific genomic loci. Instead of using publicly available ChIP data to identify sensitive targets, we selected four target genes that were among the largest responders to dCas9-mediated repression of NMNAT1 in the bulk RNA-seq experiment: DRAM1, GSK3B, TSPAN11, and ZFHX2. This approach was better suited to identify direct gene regulatory responses to metabolite production since many of the genes identified in ChIP analyses depend on metabolic enzymes interacting with transcription factors to drive genomic localization. By taking this approach, we can directly test the transcriptional effect of localization independent of transcription factor interactions. Additionally, DRAM1 and GSK3B were responsive to upregulation of dCas9-ACSS2, whereas TSPAN11 and ZFHX2 were not. We designed 3 x gRNA to localize CRISPRm 150-400 bp upstream from the TSS. RT-qPCR of target genes from RNA isolates of HEK293T cells co-transfected with target gRNA plasmid and dCas9-NMNAT1 revealed decreased expression of all targets when targeted to gene promoters (Fig. 5A-D). Interestingly, the addition of DRAM1 and GSK3B gRNA exhibited greater repression of transcription than dCas9-NMNAT1 with control gRNA (p < 0.0001, p = 0.0280), suggesting targeting of metabolic enzymes to gene promoters accentuates dCas9-NMNAT1’s transcriptional effect. Similar studies with dCas9-ACSS2 and TSPAN11 gRNA co-transfection revealed a similar phenomenon, exemplified by a relative increase in expression when dCas9-ACSS2 is targeted to the TSPAN11 promoters (p = 0.0090) (Fig. 5H). Furthermore, TSPAN11 was not identified as a DE gene in the dCas9-ACSS2 condition, indicating that locus-specific targeting of metabolic enzymes can regulate genes that are otherwise insensitive to nuclear localization and untargeted metabolite production. Taken collectively, this data positions CRISPRm as a novel tool for site-specific metabolite production and directly shows localization of metabolic enzymes as a mechanism to modulate gene expression.

**Figure 5:**
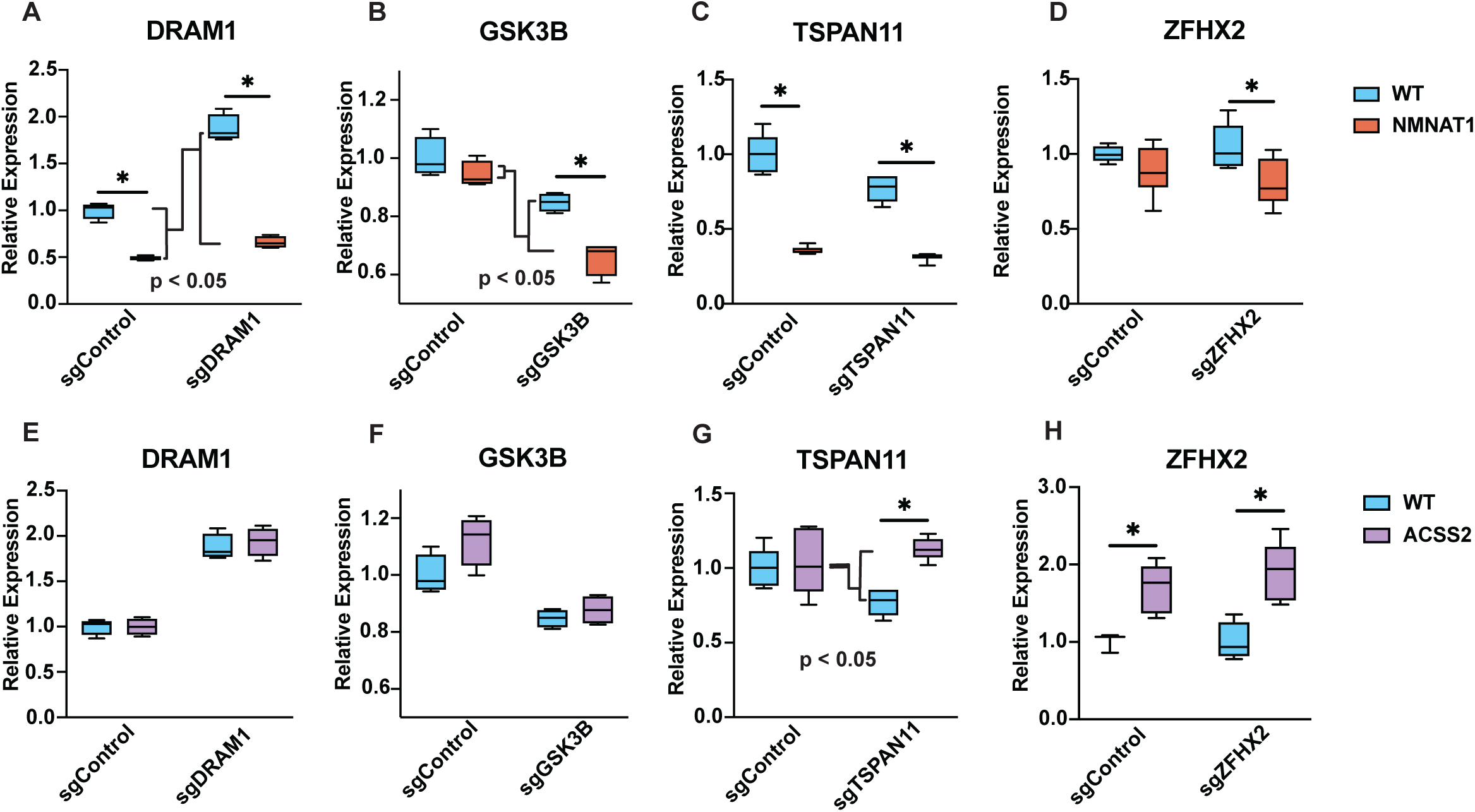
Locus-specific gene expression following targeted CRISPRm NMNAT1 and ACSS2. HEK293T cells were co-transfected with the indicated dCas9-enzyme fusion (or WT-dCas9 control) and a gene-specific gRNA targeting the promoter of DRAM1, GSK3B, TSPAN11, or ZFHX2, or a CTRL gRNA. RNA was harvested 48 hr post-transfection and analyzed by RT-qPCR. Relative expression was calculated using the ΔΔCq method, normalized to ACTB, and expressed as relative expression vs. the WT-dCas9 + non-targeting (sgCTRL) gRNA condition. (A-D) Co-transfection of dCas9-NMNAT1 with gene-specific gRNAs targeting (A) DRAM1, (B) GSK3B, (C) TSPAN11, and (D) ZFHX2. Boxplots shown; n ≥ 4 biological replicates per condition. * p < 0.05 vs. WT-dCas9 + sgControl by Student’s two-tailed t-test. (E-F) Co-transfection of dCas9-ACSS2 with gene-specific gRNAs targeting (E) DRAM1, (F) GSK3B, (G) TSPAN11, and (H) ZFHX2. Boxplots shown; n ≥ 4 biological replicates per condition. * p < 0.05 vs. WT-dCas9 + sgControl by Student’s two-tailed t-test. Two-way ANOVA was used to assess the statistical significance of the relative change in expression with dCas9-NMNAT1 or - ACSS2 compared to WT-dCas9 when targeted to each genetic locus; significance is denoted with “p < 0.05.”

## Discussion

Studies linking metabolism to chromatin state have been limited by an inability to delineate nuclear vs. cytoplasmic metabolite perturbations as drivers of transcriptional regulation. Here, we developed CRISPRm, a dCas9-metabolic enzyme fusion platform that enables nuclear and genomic localization of metabolic enzymes. CRISPRm overcomes these constraints by forcing nuclear localization of metabolic enzymes and enabling direct interrogation of whether nuclear metabolite production is sufficient to alter transcriptional output. Moreover, CRISPRm permits the directed localization of metabolic enzymes to specific gene promoters. Expression of five dCas9-enzyme fusions: dCas9-ACSS2, -NMNAT1, -MAT2A, -AHCY, and -GDH produced distinct transcriptional programs, with dCas9-ACSS2 and -NMNAT1 generating the largest transcriptional responses. Furthermore, we identified chromatin modifications and effectors enriched at shared genes that are reciprocally regulated by dCas9-ACSS2 and -NMNAT1. These findings establish CRISPRm as a novel tool for studying the function of nuclear metabolic enzymes in histone modification and gene regulation.

CRISPRm enabled the identification of candidate genes and transcriptional programs reciprocally regulated by acetyl-CoA and NAD^+^ nuclear production. dCas9-ACSS2 and -NMNAT1 expression yielded reciprocal enrichment across 68 of 70 shared GO:BP terms, which were primarily related to cell cycle regulation and ribosome assembly. Nuclear enrichment of dCas9-ACSS2 and the associated changes in acetyl-CoA support production of acetyl-CoA within the nucleus. Although dCas9-NMNAT1 expression did not significantly alter total cell NAD^+^ levels, prior studies have reported discrepancies between whole-cell and compartment-level NAD+ measurements. For instance, NAD^+^ can be specifically enriched in mitochondria without perturbing the cytosol, and decreased nuclear NAD^+^ levels are not detected by whole-cell measurement (42, 43). Since the nucleus contains only 25% of the free NAD^+^ (44), NAD^+^ would need to increase by 5-fold for a 2-fold enrichment to be detected by whole-cell LC-MS. Further studies using subcellular LC-MS metabolite quantification methods are needed to determine compartment-specific perturbation. The robust transcriptional repression of dCas9-NMNAT1 is consistent with nuclear NAD^+^ increase below the detection limit of whole-cell metabolomics. Moreover, 66% of the shared DE genes between dCas9-ACSS2 and -NMNAT1 are reciprocally regulated, suggesting that (de)acetylation mechanisms involving acetyltransferases and NAD^+^-dependent deacetylases are controlled by co-substrate levels. Potential (de)acetylation targets that could mediate the transcriptional response to nuclear acetyl-CoA and NAD+ include histones and non-histone proteins, such as transcription factors. E2F1 is a transcription factor that connects acetylation dynamics to cell cycle control. Many of the shared cell-cycle term core-enrichment genes are E2F1 targets, including MDM2, EZH2, RAD21, RB1, and AURKA. Previous studies have shown that p300/CBP acetylation promotes, and SIRT1 deacetylation opposes E2F1’s transcriptional activity (45, 46). Similarly, reciprocal acetylation/d deacetylation of Ifh1 by GCN5 and sirtuin family enzymes regulates ribosomal protein gene transcription (33). Collectively, the concordance between the reciprocally enriched pathways and known acetylation-dependent gene targets supports that the transcriptional responses are driven, at least in part, by alterations in acetylation and deacetylation activity of chromatin-modifying enzymes.

Despite the transcriptional reciprocity exhibited by dCas9-ACSS2 and -NMNAT1, there was no significant global change in histone peptide modifications. The lack of global changes in histone acetylation suggests that the relationship between nuclear acetyl-CoA and NAD^+^ pools and histone post-translational modifications is considerably more targeted than would be predicted by compartmental metabolite concentrations as the sole driver of modification state. A model of targeted changes in histone modification at specific loci is consistent with previous work describing ACSS2 localizing to specific sites and, in cooperation with HDACs and HATs, recycling stored acetate from other histone acetylation sites to sites of gene-activating acetylation (47). This model is further supported by the differential enrichment of H3K9ac, H3K18ac, H3K27ac, EP300 at ‘acetyl-sensitive’ gene promoters in public ChIP-seq analyses. Promoter-resident p300 could serve as a metabolically responsive switch: under conditions of acetyl-CoA surplus, acetylation of p300’s autoinhibitory loop relieves catalytic repression, thereby sustaining histone acetylation at target gene promoters. Conversely, under conditions of elevated NAD⁺ availability, deacetylation of the autoinhibitory loop renders p300 catalytically inactive and attenuates acetylation of nearby loci. Further work is needed to determine the role of loci-specific histone acetylation changes and the histone-modifying enzymes involved. Notably, the observation that certain loci remained transcriptionally unresponsive to global increases in nuclear acetyl-CoA mediated by untargeted ACSS2 expression, yet exhibited significant upregulation upon direct promoter targeting of the same enzyme, implies the existence of locus-specific barriers that preclude a subset of genomic regions from responding simply to nuclear metabolite levels. The capacity of promoter-targeted CRISPRm to overcome these barriers suggests that forced proximity of metabolic enzyme activity to chromatin substrates can bypass such constraints, providing evidence that the spatial organization of nuclear metabolism is a critical determinant of epigenetic responsiveness.

CRISPRm also identified candidate genes and transcriptional programs sensitive to methylation. dCas9-MAT2A displayed a robust activating gene signature, and trending enrichment of H3K4me3 and H3K27me3K36me2 consistent with prior reports of SAM-driven H3K4me3 and H3K36me2 (48). Prior studies have shown that KDM2A/B demethylation of H3K36me2 and non-histone targets are associated with diverse gene regulatory programs (49–51). dCas9-GDH displayed a primarily repressive transcriptional signature and trends toward reductions in H3K36me2, consistent with increased αKG-dependent demethylase activity at activating marks despite the absence of whole-cell αKG changes. This apparent discrepancy may reflect the limitations of steady-state metabolite quantification, which does not account for the rapid flux of αKG through downstream pathways. In fact, one metabolic flux analysis inferred the net flux through GDH is nearly 50 mM/hr (52), highlighting the rapid replacement of αKG levels upon use by downstream enzymes. Further work using isotopic labeling, and flux analyses will be required to assess downstream turnover of αKG.

dCas9-AHCY was predominantly repressive. AHCY has been shown to increase DNA methylation in HEK293 cells and maintain repressive H3K27me3 at developmental gene loci (53, 54). Interestingly, dCas9-MAT2A and-AHCY, which are thought to promote methylation via distinct mechanisms, exhibited largely opposite enrichment in the shared GO:BP terms identified. AHCY consumes SAH, a product inhibitor of SAM-dependent methyltransferases, potentially relieving feedback inhibition. The results presented here suggest that reduced levels of SAH favor a more repressive chromatin state where the balance of opposing methyltransferases would suggest that H3K4 or H3K36 methyltransferases are less sensitive to SAH product inhibition and that PRC2 catalyzed methylation of K27 is enhanced upon SAH reduction. This differential sensitivity would be expected to yield a net shift in the histone methylation landscape, favoring repressive marks at the expense of transcriptionally activating modifications. Consistent with the observed overall decrease in transcriptional output with dCas9-AHCY expression, the data showed trends of decreased K36 methylation within H3K27me3K36me2. The fact that dCas9-MAT2A led to an opposing gene up-regulation suggests that H3K4 or H3K36 methyltransferase respond more to increased SAM levels than reduced SAH levels. Thus, these methyltransferases are not simply regulated by the ratio of SAM/SAH but also by the enzymes’ intrinsic catalytic properties (65). The relative K_m_ values for SAM and the inhibition constant K_i_ for SAH must be considered to explain the sensitivity to varied levels of SAM and SAH. Further work will be needed to establish whether these catalytic differences can explain the cellular observations. Our mechanistic analyses focused on histone modifications as a mediator of observed transcriptional regulation, but metabolites influence gene regulation through alternative mechanisms. DNA and RNA methylation are also sensitive to perturbations in SAM and SAH availability and likely contribute to the transcriptional phenotypes observed here. Future studies examining the impact of nuclear methyl metabolite production on DNA and RNA methylation will be necessary to delineate the relative contributions of histone versus nucleic acid methylation to the transcriptional changes observed here.

In addition to demonstrating that CRISPRm can be used to assess regulatory mechanisms that respond to fluctuations in nuclear metabolites, these studies provide evidence that targeting metabolic enzymes to promoters is sufficient to potentiate the transcriptional response to that enzyme compared to when localized to the nucleus. dCas9-NMNAT1 exhibited enhanced transcriptional repression upon promoter-directed targeting, while dCas9-ACSS2 induced upregulation at loci previously characterized as insensitive in our untargeted RNA-seq analyses. These observations suggest that localized metabolite production at specific genomic loci is a viable mechanism by which chromatin-modifying substrates are enriched at defined regulatory elements, thereby conferring post-translational modifications that drive gene regulation. This principle is well-established in the context of metabolons, multienzyme complexes that facilitate metabolite channeling among members to increase local reaction rates and shield intermediates from competing pathways. The SESAME complex, for instance, functions as a chromatin-associated metabolon that senses cellular metabolic state and locally generates SAM to regulate histone methylation and transcriptional output (55). While we focused on the role of nuclear perturbation by wild-type enzymes, the metabolic enzymes employed as dCas9 fusion partners could possess non-catalytic functions in transcriptional regulation through direct protein–protein interactions or recruitment of regulatory complexes, as has been described for ACSS2 and NMNAT1 (39, 56, 57). Future work is needed to identify the relative contribution of catalytic and non-catalytic mechanisms of gene regulation. Moreover, whether CRISPRm can exhibit similar phenotypes when targeted to other cis-regulatory elements, such as silencers or enhancers, remains to be studied.

In conclusion, CRISPRm is a novel tool for perturbing nuclear metabolites relevant to chromatin-modifying enzymes. This platform enables the study of chromatin responses to metabolite perturbation in the nucleus and at specific genomic loci. Here, we applied the CRISPRm platform to identify candidate acetylation- and methylation-sensitive genes and gene regulatory programs. Moreover, CRISPRm helped identify candidate chromatin features that confer transcriptional responsiveness to nuclear metabolite perturbation. More broadly, CRISPRm can identify spatial mechanisms of gene regulation, including the localization of metabolic enzymes to promoters, to further our understanding of metabolism-chromatin crosstalk.

## Materials and Methods

### In vitro Cell Culture

*H. sapiens* HEK293T cell lines were cultured in RPMI 1640 media (Gibco™, 11875119) and 10% FBS (Gibco™, A5670701) at 37°C, 5% CO_2_.

### DNA Transfections

HEK293T cells were seeded 24 hours prior to DNA transfection, with RPMI culture media being replaced 1 hour prior to the addition of prepared lipoprotein particles. All DNA stocks used for transfections were generated using the PureYield™ Plasmid Midiprep System (Promega, A2492). DNA plasmids were transfected using a 4:1 reagent:DNA ratio using FuGENE® HD transfection reagent (Promega, E2311). Input DNA mass was calculated for the transfection of equimolar concentrations across conditions for no sgRNA experiments (i.e., HA-dCas9 only) and between HA-dCas9 and sgRNA plasmids for co-transfection experiments. Cells were cultured for 48 hours before harvesting for downstream analyses.

### CRISPR Target Sequence Design

To identify CRISPR target sequences, the hg38 assembly for each gene of interest was accessed via the National Center for Biotechnology Information (NCBI) webpage after which the depicted range was adjusted to isolate 500 bp of sequence upstream of the corresponding transcription start site (TSS). This sequence was analyzed using CHOPCHOP (https://chopchop.cbu.uib.no, PMID: 31106371) to generate a ranked list of CRISPR target sequences of which three were chosen for use in downstream assays based on additional criteria including a requirement the sequence is located within -75 bp to -450 bp upstream of the TSS and a lack of target sequence overlap (58). An additional guanosine nucleotide was added to CRISPR target sequences as needed to maximize each individual sgRNAs transcription efficiency. The control sgRNA sequence was the only sequence not generated as described above and was instead taken from Martin et al., 2018 (59).

### sgRNA Plasmid Engineering

To generate an sgRNA plasmid capable of expressing three individual sgRNAs, the plasmid pX333 was first acquired as a gift from Andrea Ventura (Addgene plasmid # 64073 ; http://n2t.net/addgene:64073 ; RRID:Addgene_64073) (60). pX333, which is capable of expressing two individual sgRNAs, was restriction digested at the KpnI sequence (Thermo Scientific™, FD0524) followed by insertion of a third U6 promoter and gRNA scaffold via NEBuilder® HiFi DNA Assembly (New England Biolabs®, E2621S). This third gRNA scaffold incorporated the SapI Type IIs target sequence to facilitate sgRNA insertion. The engineered 3x sgRNA expression plasmid (i.e., pX336) was then restriction digested at the EcoRI sequence (Thermo Scientific™, FD0274) for insertion of a T2A-eGFP cassette at the dCas9 C-terminus via NEBuilder® HiFi DNA Assembly (i.e., pX336-Cas9-eGFP) (New England Biolabs®, E2621S). Finally, the entire Cas9-eGFP cassette was removed via AgeI/EcoRI double restriction digest (Thermo Scientific™, FD1464 and FD0524) followed by NEBuilder® HiFi DNA Assembly (New England Biolabs®, E2621S) insertion of a stand-alone eGFP cassette, generating the final pX336-NoCas9-eGFP plasmid.

To insert sgRNAs into pX336-NoCas9-eGFP, individual sgRNA primer pairs were designed with appropriate overlap for downstream T4 ligation into the first, second, or third sgRNA scaffolds as needed. Individual sgRNA primers were reconstituted to a final concentration of 100μM in Duplex Buffer (i.e., 100mM potassium acetate, 30mM HEPES, pH 7.5). Paired forward and reverse primers of 1 uL each were added to 18μl of Duplex Buffer and annealed on a Veriti™ Thermal Cycler with the following protocol steps: Stage 1 – 5 min at 95°C, Stage 2 –10 min at 25°C with a 3% ramp rate, Stage 3 – infinite hold at 4°C. Annealed primers were diluted 1:125 in Duplex Buffer and stored at -20°C. Next, the pX336-NoCas9-eGFP backbone was restriction digested at the BbsI sequence (Thermo Scientific™, FD1014), followed by additional cleanup using the QIAquick PCR Purification Kit (Qiagen, 28104). Next, the appropriate sgRNA duplex was inserted into the digested backbone using the NEB T4 DNA Ligase (New England Biolabs®, M0202S). T4 ligation reaction was transformed into Stellar Chemically Competent *E. coli* (Takara, 636763) followed by plasmid isolation using the GeneJET Plasmid Miniprep Kit (Thermo Scientific™, K0503). This process was repeated sequentially for the insertion of sgRNAs two and three, with the sequence of all intermediate and final plasmids being confirmed via Plasmidsaurus whole-plasmid sequencing.

### CRISPRm dCas9 Construct Engineering

All dCas9 constructs were engineered based on the previously published dCas9-KRAB which was a gift from Alejandro Chavez & George Church (Addgene plasmid # 110820 ; http://n2t.net/addgene:110820 ; RRID:Addgene_110820) (28). First, an SV40 promoter driven mCherry cassette was inserted after the HSV TK poly(A) signal via via NEBuilder® HiFi DNA Assembly (New England Biolabs®, E2621S) using dCas9-KRAB backbone and mCherry cassette PCR fragments. Next, the dCas9-KRAB-mCherry plasmid was restriction digested at the XbaI sequence (Thermo Scientific™, FD0684) for insertion of a 2X HA epitope tag onto the dCas9 N-terminus via NEBuilder® HiFi DNA Assembly (New England Biolabs®, E2621S). The 2X HA epitope was designed and amplified off an Integrated DNA Technologies (IDT) gBlock™. Finally, the C-terminal KRAB domain and linker sequence was removed via backbone PCR amplification followed by T4 DNA ligase blunt end ligation, leaving a final HA-dCas9-mCherry construct.

To insert metabolic enzymes onto the HA-dCas9 C-terminus, backbone constructs were restriction digested at the EcoRV sequence (Thermo Scientific™, FD0303) which immediately followed the HA-dCas9 stop codon. Metabolic enzymes were then PCR amplified off of (1) IDT gBlocks™ for ACSS2, AHCY, GDH, (2) pMXs_FLAG-NMNAT1 which was a gift from David Sabatini (Addgene plasmid # 133259 ; http://n2t.net/addgene:133259 ; RRID:Addgene_133259) for NMNAT1, or (3) a Denu laboratory legacy plasmid for MAT2A (61). All amplified DNA sequences encoded the canonical, *H. sapiens* UniProt amino acid sequences except for GDH due to the intentional omission of the protein’s mitochondrial targeting signal. All amplified DNA fragments were inserted into the HA-dCas9-mCherry backbone via NEBuilder® HiFi DNA Assembly (New England Biolabs®, E2621S). The sequence of all intermediate and final plasmids were confirmed via Plasmidsaurus whole-plasmid sequencing.

### Immunoblotting

Tissue culture samples were lysed in ice cold RIPA buffer supplemented with 10 μg/ml leupeptin, 10 μg/ml aprotinin, 100 μM phenylmethylsulfonyl fluoride protease inhibitors for 30 min on ice with additional vortexing every 10 min. Next, samples were centrifuged at 14,000xg for 20 min at 4°C after which the supernatant was transferred to a new 1.5 ml Eppendorf tube. Protein concentrations for each sample were quantified using Pierce BCA Protein Assay reagents, with 30 μg of protein being separated by SDS-PAGE on 7.5% resolving gels before subsequent transfer to 0.2 µm pore nitrocellulose membrane at 20V for 24 hours. Membranes were blocked in 0.1% fat free milk PBS for 1 hour followed by overnight primary antibody incubation (anti-HA: EpiCypher, #13-2010, 1:1000, host – rabbit; anti-alpha Tubulin, Cell Signaling Technology®, #3873S, 1:1000, host – mouse) and 1 hour of secondary antibody incubation (IRDye® 800CW anti-Rabbit: LICORbio™, #926-32211, 1:10,000, host – goat; IRDye® 680RD anit Mouse: LICORbio™, #925-68070, 1:10,000, host – goat) in 5% fat free milk PBST. Membranes were imaged using an Odyssey Infrared Imager (model no. 9120).

### Metabolite Extraction

HEK293T cells were washed 2x with 1 ml of ice-cold PBS pH 7.4 and incubated in 1 ml of -80 °C 40:40:20 MeOH:ACN:H_2_O extraction solvent for 15 min at -80 °C. Cells were scraped and transferred to a 1.5 ml Eppendorf tube followed by 5 sec of vortexing and a 5 min incubation on ice. Samples were then centrifuged at 21,100xg for 5 min at 4°C, after which the supernatant was transferred to a new 1.5 ml Eppendorf tube and dried using a Thermo Fisher Savant ISS110 SpeedVac. The remaining cell pellet was used for RIPA protein extraction and BCA protein quantification to generate normalization factors for LC-MS metabolomics data analyses. Finally, dried metabolite extracts were resuspended in 175 µl of H_2_O for hydrophobic compound detection or 150 µl of 80% ACN with 20% H_2_O (20mM final ammonium acetate and 20mM final ammonium hydroxide) for polar compound detection and centrifuged at 21,100xg for 5 min at 4°C with supernatants being transferred to glass vials for LC-MS analysis.

### LC-MS Metabolomics

Metabolite samples resuspended in H_2_O were injected in random order onto a Thermo Fisher Scientific Vanquish UHPLC with a Waters Acquity UPLC BEH C18 column (1.7 μm, 2.1 x 100mm; Waters Corp., Milford, MA, USA) and analyzed using a Thermo Fisher Q Exactive Orbitrap mass spectrometer in negative ionization mode. LC separation was performed over a 25 minute method with a 16 minute step-wise gradient of mobile phase (buffer A, 97% water with 3% methanol, 10mM tributylamine, and acetic acid-adjusted pH of 8.3) and organic phase (buffer B, 100% methanol) (0 minute, 5% B; 2.5 minute, 5% B; 5 minute, 20% B; 7.5 minute, 20% B; 13 minute, 55% B; 15.5 minute, 95% B; 18.5 minute, 95% B; 19 minute, 5% B; 25 minute, 5% B, flow rate 0.2mL/min). A quantity of 15 μl of each sample was injected into the system for analysis. The ESI settings were 30/10/1 for sheath/aux/sweep gas flow rates, 2.50kV for spray voltage, 50 for S-lens RF level, 350°C for capillary temperature, and 300°C for auxiliary gas heater temperature. MS1 scans were operated at resolution = 70,000, scan range = 85-1250m/z, automatic gain control target = 1 x 10^6^, and 100 ms maximum IT.

Metabolite samples resuspended in 80% ACN with 20% H_2_O (20mM final ammonium acetate and 20mM final ammonium hydroxide) were injected in random order onto a Thermo Fisher Scientific Vanquish UHPLC with a Waters XBridge BEH Amide column (130Å, 2.5 μm, 2.1 x 150 mm; Waters Corp., Milford, MA, USA) and analyzed using a Thermo Fisher Q Exactive Orbitrap mass spectrometer in positive ionization mode. LC separation was performed over a 22 minute method with a 13 minute step-wise gradient of mobile phase (buffer A, 5% ACN with 95% methanol, 20 mM ammonium acetate and 20mM ammonium hydroxide) and organic phase (buffer B, 80% ACN with 20% H_2_O, 20 mM ammonium acetate and 20mM ammonium hydroxide) (0 minute, 100% B; 3 minute, 100% B; 3.2 minute, 90% B; 6.2 minute, 90% B; 6.5 minute, 80% B; 10.5 minute, 80% B; 10.7 minute, 70% B; 13.5 minute, 70% B; 13.7 minute, 45% B; 16 minute, 45% B; 16.5 minute, 100% B; 22 minute, 100% B; flow rate 0.3 mL/min). A quantity of 15 μl of each sample was injected into the system for analysis. The ESI settings were 10/5/1 for sheath/aux/sweep gas flow rates, 3.50kV for spray voltage, 50 for S-lens RF level, 350°C for capillary temperature, and 30°C for auxiliary gas heater temperature. MS1 scans were operated at resolution = 70,000, scan range = 60-186m/z and 187-900m/z, automatic gain control target = 3 x 10^6^, and 200 ms maximum IT.

Raw data files were converted into mzml for metabolite identification and peak AreaTop quantification using El-MAVEN (v0.12.1-beta) (62).

### Histone Protein Acid Extraction and Derivatization

Flash frozen HEK293T cell pellets were thawed and resuspended in 800 μl of ice-cold Buffer A (10 mM Tris-HCl pH 7.4, 10 mM NaCl, and 3 mM MgCl2) supplied with protease and histone deacetylase inhibitors (10 μg/ml leupeptin, 10 μg/ml aprotinin, 100 μM phenylmethylsulfonyl fluoride, 10 mM nicotinamide, and 4 μM trichostatin A) followed by 80 strokes of loose--pestle homogenization in a 1 mLlWheaton dounce homogenizer before being transferred to a new 1.5 ml Eppendorf tube. Samples were then centrifuged at 800xg for 10 minutes at 4°C to pellet nuclei. Supernatants were discarded and the remaining nuclei pellet was resuspended in 500 μl ice-cold PBS pH 7.4 and centrifuged at 800xg for 10 minutes at 4°C. The supernatant was discarded and nuclei were again washed as previously described. Next, pelleted nuclei were resuspended in 500 μl of 0.4N H_2_SO_4_ and rotated at 4°C for 4 hours. Samples were centrifuged at 3,400xg for 5 minutes at 4°C to pellet nuclear debris and precipitated non-histone proteins. Supernatants were transferred to new 1.5 ml Eppendorf tubes after which 125 μl of 100% trichloroacetic acid was added and incubated overnight on ice at 4°C. The next day, samples were centrifuged at 3,400xg for 5 minutes at 4°C to pellet precipitated histone proteins. Supernatants were discarded and the precipitant was washed with 1 ml of ice-cold acetone +0.1% HCl. Samples were centrifuged at 3,400xg for 2 minutes at 4°C and supernatants were again discarded. This process was repeated with a 100% ice-cold acetone wash. Residual acetone was allowed to evaporate at room temperature for 10 minutes after which dried histone protein precipitant was resuspended in 100 μl H_2_O. Finally, samples were centrifuged at 21,100xg for 5 minutes at 4°C to pellet any remaining insoluble debris and supernatants containing purified histone proteins were transferred to new 1.5 mL Eppendorf tubes and stored at -20°C prior to additional downstream processing.

To prepare purified histone samples for LC-MS/MS analysis, 5 μg of each sample was diluted with H_2_O to a final volume of 10 μl. 1 μl of 1 M triethylammonium bicarbonate (TEAB) was added to each sample , bringing the final pH within a range of pH 7-9. Next, 1 μl of 1:50 d6-acetic anhydride:H_2_O was added to each sample followed by a 2-minute room temperature incubation. The reaction was quenched via addition of 1 μl 80 mM hydroxylamine followed by a 20-minute room temperature incubation. Next, d3-acetylated histones were digested with 0.1 μg trypsin overnight hours at 37°C. Upon completion of trypsin digestion, prepared histone peptides were N-terminally modified with 1 μl 1:50 phenyl isocyanate:ACN for 1-hour at 37°C. Modified peptides were desalted and eluted-off of EmporeC18 extraction membrane. Eluted samples were dried completely using a Thermo Fisher Scientific Savant ISS110 SpeedVac and resuspended in 30 μl sample diluent (94.9% H2O, 5% ACN, 0.1% TFA). Resuspended samples were centrifuged at max speed for 5 min at 4°C after which supernatants were transferred to glass vials for LC-MS/MS analysis.

### LC-MS/MS Histone Proteomics

Derivatized histone peptides were injected using a Dionex Ultimate3000 nanoflow HPLC with a Waters nanoEase UPLC C18 column (100 m x 150 mm, 3 μm) onto a Thermo Fisher Eclipse mass spectrometer in positive ionization mode. LC separation was performed over a 90-minute method with a 60-minute linear gradient of mobile phase (buffer A, H_2_O + 0.1% formic acid) and organic phase (buffer B, ACN + 0.1% formic acid) (0 min, 2.0% B; 5 min, 2.0% B; 65 min, 35% B; 67 min, 95% B; 77 min, 95% B; 79 min, 2.0% B; 90 min, 2.0% B). A quantity of 3 μl of each sample was injected into the system for analysis. MS1 scans were operated at resolution = 60,000, scan range = 395-1005 m/z, automatic gain control target = 1 x 10^6^, and 100 ms maximum IT followed by a DIA scan with a loop count of 30. DIA scans were operated at window size = 20 m/z, resolution = 15,000, automatic gain control target = 1 × 10^5^, HCD collision energy = 30 and 40 ms maximum IT.

DIA Thermo .raw files were analyzed via the EpiProfile 2.0 AcD3 module with a mass tolerance of 8 ppm (63). Subsequent data filtering (i.e., removing samples with (1) >2 null values for common peptides or (2) >50% CV) and normalization was performed using our published Histone Analysis Workflow (64). Heatmaps of histone peptide log fold changes were constructed using R package pheatmap (v1.0.13).

### RNA Isolation and cDNA Synthesis

Total RNA was isolated from frozen HEK293T cell pellets using the Monarch® Spin RNA Isolation Kit (New England Biolabs®, T2110S), including optional steps for genomic DNA digestion. Next, the RevertAid First Strand cDNA Synthesis Kit (Thermo Scientific™, K1622) was used to generate cDNA from 750 ng of isolated RNA with random hexamer primers. Newly synthesized cDNA was diluted 1:3 in H_2_O and stored at -20°C.

### RT-qPCR

To quantitatively measure target gene expression, 2 μl of prepared cDNA was used as input material for PowerUp™ SYBR™ Green Master Mix (Applied Biosystems™, A25742) qPCR reactions. All qPCR reactions were analyzed using a Bio-Rad CFX Opus 96 Real-Time PCR System an internally normalized using beta-Actin abundance. All primer pair sequences used in qPCR reactions are provided in Table S1. Novel primer pair sequences generated for this study were obtained using the NCBI Primer-Blast tool with default settings except (1) an adjusted product size range of 70 – 200 bp and (2) allowing for primers to amplify mRNA splice variants. Relative mRNA expression was calculated by the 2^-ΔΔCt method with ACTB as the reference gene.

### RNA-sequencing and Data Analysis

Isolated, total RNA samples were provided to Azenta Life Sciences for Poly(A) selection/library preparation and Illumina 2x150 bp sequencing with a target sequencing depth of ∼30 million reads per sample. Adapters were trimmed from raw .fastq sequencing files using trimmomatic (v0.39) followed by transcript alignment and quantification using RSEM (v1.3.3) with the publicly available hg18 human indices. RSEM output .gene files were subsequently processed in RStudio (v2025.05.0+496) to extract expected count values for pairwise differential gene expression analysis via DESeq2 (v1.48.1). Gene Set Enrichment Analysis of total genes ranked by Wald statistic was performed in Rstudio using clusterProfiler gseGO command (v4.16.0) with the following settings: OrgDb = org.Hs.eg.db; keyType = "SYMBOL"; ont = "BP”; pvalueCutoff = 0.05; pAdjustMethod = "BH"; seed = TRUE. Differentially expressed genes were filtered based on a padj < 0.05 and a 0.25 log2FC cutoff. Overrepresentation analysis of acyl- or methyl-sensitive DE genes was performed in RStudio using the clusterProfiler enrichGO command (v4.16.0) with the following settings: OrgDb = org.Hs.eg.db; keyType = "SYMBOL"; ont = "BP"; pAdjustMethod = "BH"; pvalueCutoff = 1; minGSSize = 10; universe used was all identified genes in the DESeq output.

### Analysis of Publicly Available ChIP-Seq Data

BigWig files from ChIP-seq in HEK293T cells were downloaded from Gene Expression Omnibus as follows:

| Genome used for alignment | GEO Sample | PTM |
| --- | --- | --- |
| hg18 | GSM1249885 | H3K4me3 |
| hg18 | GSM1249890 | P300 |
| hg19 | GSM6249166 | H3K9ac |
| hg19 | GSE81696 | H3K14ac |
| hg19 | GSE130137 | H3K18ac |
| hg |  | H3K27ac |
| hg |  | H3K4me1 |

UCSC’s bigWigLiftOver (version 482) was used to convert BigWig files originally aligned to hg18 to hg19. Deeptools’s (version 3.5.5) “computeMatrix scale-regions -b 3000 -a 3000” command was used to calculate signal enrichment 3kb up and downstream and across stretched gene bodies. Deeptools “plotHeatmap” command was used to plot heatmaps and metaplots for signal enrichment of ACSS2 upregulated genes, NMNAT1 downregulated genes, and methyl-sensitive genes (differentially expressed in dCas9-MAT2A, AHCY, or GDH compared to WT).

### Statistics

Statistical analyses were conducted using Prism, version 10 (GraphPad Software Inc.) or R (version 4.5.1) unless otherwise indicated. Peak AreaTop of LC-MS metabolomic data were compared using an unpaired two-tailed student’s t-test. Relative mRNA expression from RT-qPCR data was compared using an unpaired two-tailed Student’s t-test of ΔCt values. Histone PTM stoichiometry was compared across conditions using limma with Benjamini–Hochberg correction (v3.66.0).

### Data Availability

Bulk RNA-sequencing data has been uploaded to Gene Expression Omnibus, accession number GSE######. Raw mass spectrometry data files for the LC-MS metabolomics (MSV000101721) and histone proteomics (MSV000101748) have been uploaded to the MassIVE database.

## Acknowledgments

The Denu lab is supported in part by the NIH (GM149279 and DK125859). JMD is a member of the Wisconsin Nathan Shock Center of Excellence in the Basic Biology of Aging, P30 AG09258601. This research was supported by The New York Stem Cell Foundation, SAH is a NYSCF - Druckenmiller Fellow. All flow cytometry was performed with the assistance of the University of Wisconsin Carbone Cancer Center Flow Cytometry Laboratory (University of Wisconsin Carbone Cancer Center Support Grant P30 CA014520).

## Declaration of Competing Interests

JMD is a co-founder of Galilei Biosciences and a consultant for Evrys Bio.

## Notes

### Competing Interest Statement

JMD is cofounder of Galilei Biosciences and a consultant for Evrys Bio.

