## Supplemental Figures for "dCas9-metabolic enzyme fusions modulate global and locus-specific gene expression"

#### Figure S1: Tree Plots of CRISPRm Enriched GO:BP Terms

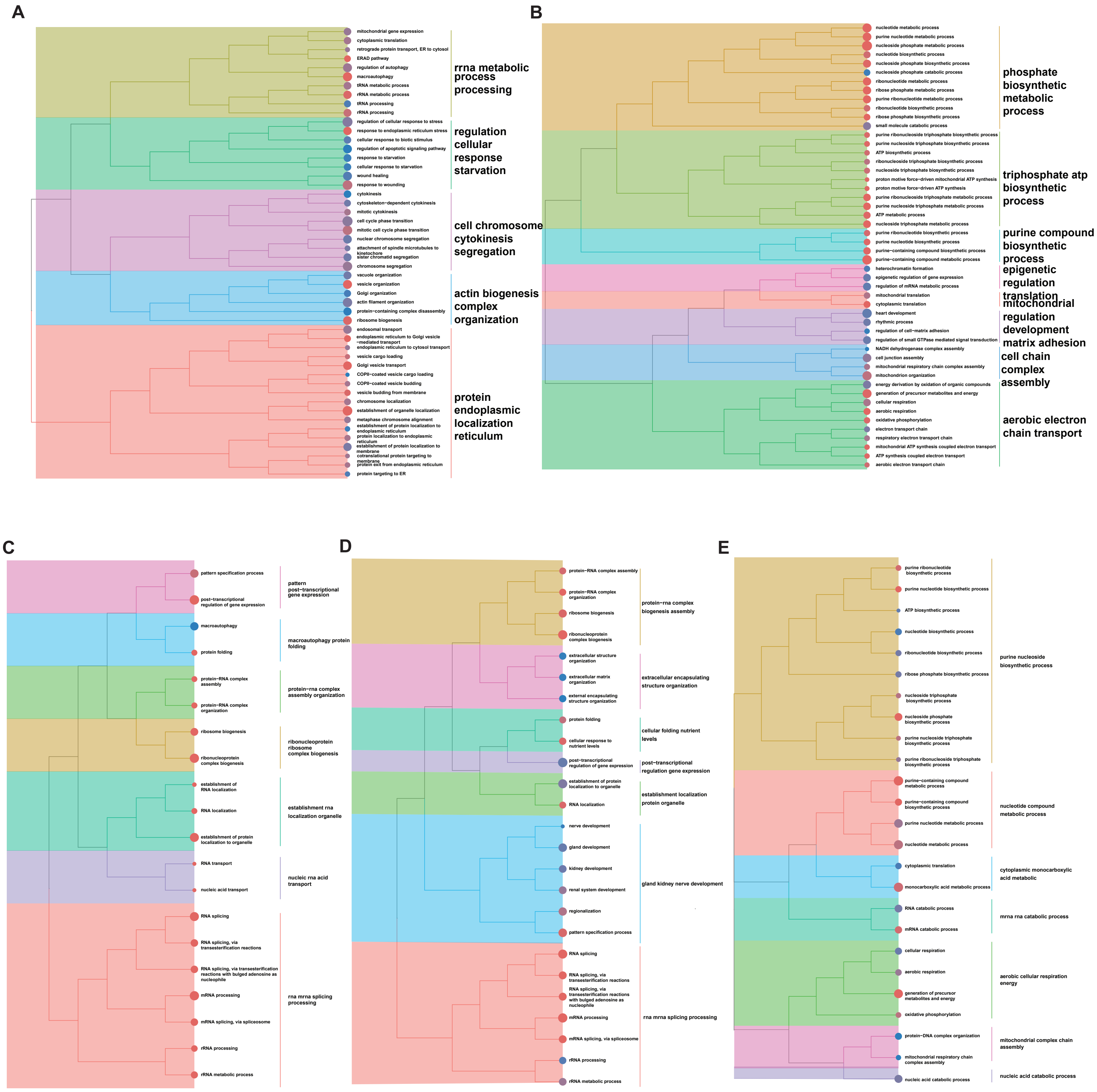


**Figure S1:** ***Tree Plots of CRISPRm RNAseq Enriched GO:BP Terms.*** (A–E) Tree plot visualizations of the top 50 GO Biological Process terms enriched by GSEA, ranked by Wald statistic, for dCas9-ACSS2 (A), -NMNAT1 (B), -MAT2A (C), -AHCY (D), and -GDH (E) relative to WT-dCas9. Semantic similarity between terms was calculated using the Wang method, and hierarchical clustering groups functionally related terms into colored modules; representative summary keywords for each module are shown at right. Plots were generated with enrichplot.

Lore ipsum

#### Figure S2: Gene set enrichment analysis of CRISPRm pertubations.

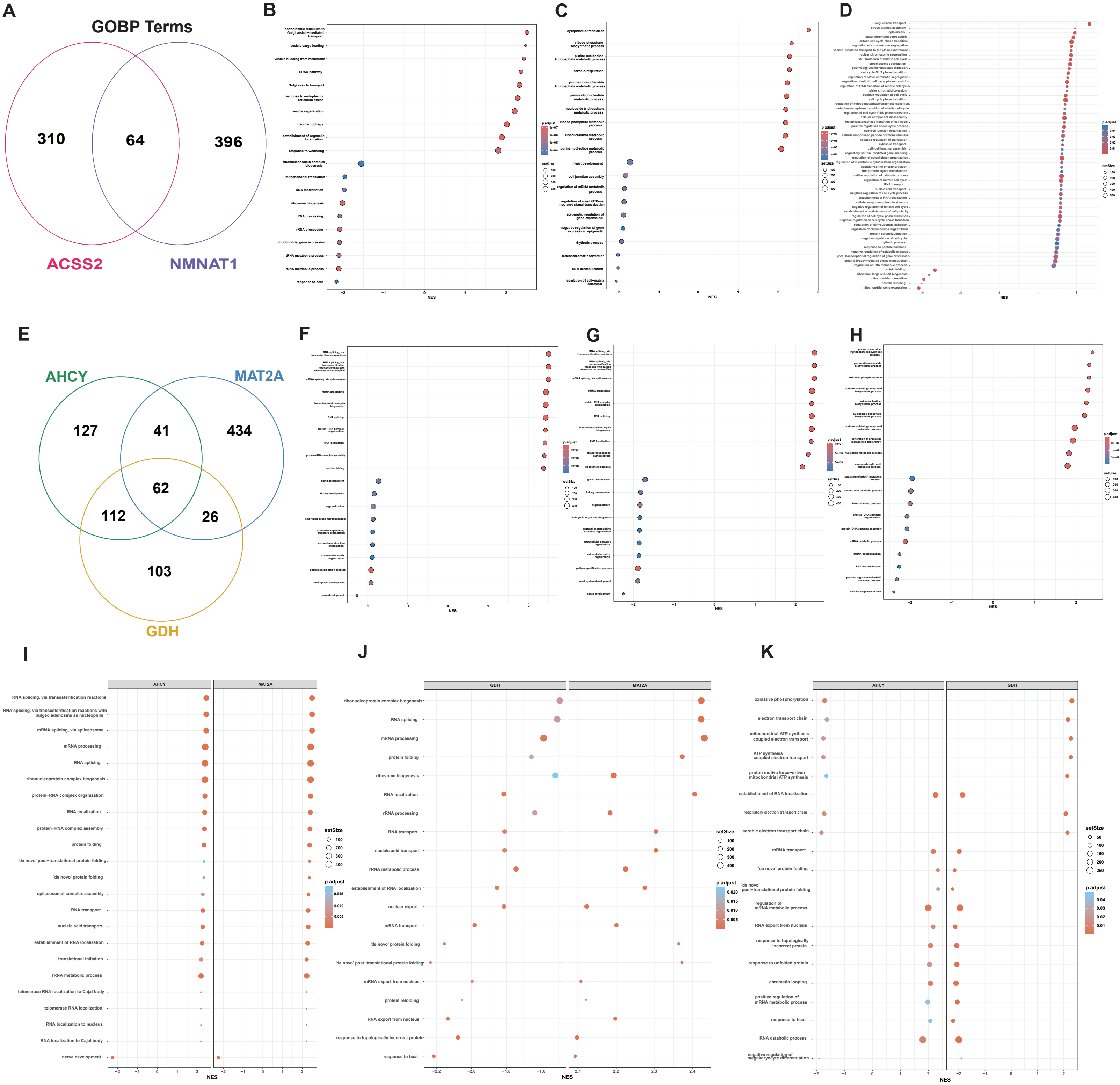


**Figure S2:** ***Gene Set Enrichment Analysis of CRISPRm Perturbations*** (A) Venn diagram showing overlap of significantly enriched GO:BP terms (padj < 0.05) identified by GSEA for dCas9-ACSS2 and -NMNAT1 relative to WT-dCas9. (B, C) Dot plots showing the top 10 up- and down-regulated GO:BP terms for (B) dCas9-ACSS2 and (C) dCas9-NMNAT1, ranked by normalized enrichment score (NES); dot color reflects Benjamini-Hochberg p-val. (D) Dot plot of the top 50 GO:BP terms shared between dCas9-ACSS2 and -NMNAT1\(E) Venn diagram showing overlap of significantly enriched GO:BP terms across dCas9-MAT2A, -AHCY, and -GDH relative to WT-dCas9. (F–H) Dot plots showing the top 10 up- and down-regulated GO:BP terms for (F) dCas9-MAT2A, (G) dCas9-AHCY, and (H) dCas9-GDH, encoded as in B and C. (I–K) Side-by-side dot plots of the top 50 GO:BP terms shared between (I) MAT2A and AHCY, (J) MAT2A and GDH, and (K) AHCY and GDH.

### Figure S3: CRISPRm Histone PTM Clustered Heatmaps

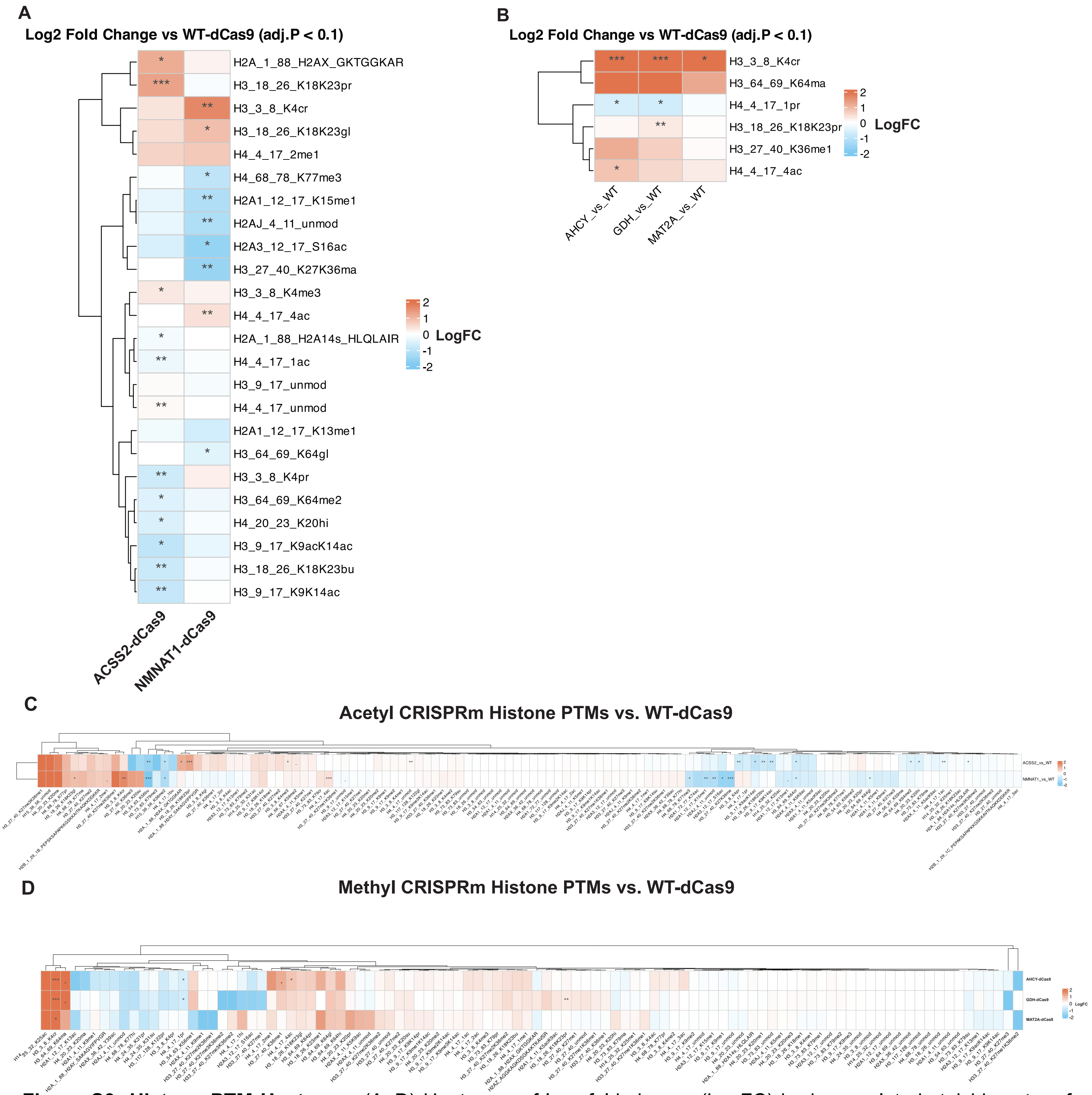


**Figure S3: Histone PTM Heatmaps** (A–D) Heatmaps of log₂ fold change (LogFC) in deconvoluted stoichiometry of residue-specific histone post-translational modifications (PTMs) for each dCas9 fusion relative to WT-dCas9. Color indicates LogFC (red, increased; blue, decreased), and asterisks denote significant changes ((., padj < 0.1; *, padj < 0.05; ***, padj < 0.01) [limma]). Rows were clustered hierarchically using [Euclidean distance and complete linkage]; columns were not clustered. (A) Acetyl PTMs, restricted to peptides with a significant change in at least one perturbation (padj < 0.1). (B) Methyl PTMs, filtered as in A. (C) All quantified acetyl PTMs, unfiltered. (D) All quantified methyl PTMs, unfiltered.
